# NetPyNE: a tool for data-driven multiscale modeling of brain circuits

**DOI:** 10.1101/461137

**Authors:** Salvador Dura-Bernal, Benjamin A Suter, Padraig Gleeson, Matteo Cantarelli, Adrian Quintana, Facundo Rodriguez, David J Kedziora, George L Chadderdon, Cliff C Kerr, Samuel A Neymotin, Robert McDougal, Michael Hines, Gordon M G Shepherd, William W Lytton

## Abstract

Biophysical modeling of neuronal networks helps to integrate and interpret rapidly growing and disparate experimental datasets at multiple scales. The NetPyNE tool (www.netpyne.org) provides both programmatic and graphical interfaces to develop data-driven multiscale network models in NEURON. NetPyNE clearly separates model parameters from implementation code. Users provide specifications at a high level via a standardized declarative language, *e.g.,* a connectivity rule, instead of tens of loops to create millions of cell-to-cell connections. Users can then generate the NEURON network, run efficiently parallelized simulations, optimize and explore network parameters through automated batch runs, and use built-in functions for visualization and analysis – connectivity matrices, voltage traces, raster plots, local field potentials, and information theoretic measures. NetPyNE also facilitates model sharing by exporting and importing using NeuroML and SONATA standardized formats. NetPyNE is already being used to teach computational neuroscience students and by modelers to investigate different brain regions and phenomena.

## 1 Introduction

The worldwide upsurge of neuroscience research through the BRAIN Initiative, Human Brain Project, and other efforts is yielding unprecedented levels of experimental findings from many different species, brain regions, scales and techniques. As highlighted in the BRAIN Initiative 2025 report,^1^ these initiatives require computational tools to consolidate and interpret the data, and translate isolated findings into an understanding of brain function. Biophysically-detailed multiscale modeling (MSM) provides a unique method for integrating, organizing and bridging these many types of data. For example, data coming from brain slices must be compared and consolidated with *in vivo* data. These data domains cannot be compared directly, but can be potentially compared through simulations that permit one to switch readily back-and-forth between slice-simulation and *in vivo* simulation. Furthermore, these multiscale models permit one to develop hypotheses about how biological mechanisms underlie brain function. The MSM approach is essential to understand how subcellular, cellular and circuit-level components of complex neural systems interact to yield neural function and behavior.^2–4^ It also provides the bridge to more compact theoretical domains, such as low-dimensional dynamics, analytic modeling and information theory.^5–7^

NEURON is the leading simulator in the domain of multiscale neuronal modeling.^8^ It has 648 models available via ModelDB,^9^ and over 2,000 NEURON-based publications (neuron.yale.edu/neuron/publications/neuron-bibliography). However, building data-driven large-scale networks and running parallel simulations in NEURON is technically challenging,^10^ requiring integration of custom frameworks needed to build and organize complex model components across multiple scales. Other key elements of the modeling workflow such as ensuring replicability, optimizing parameters and analyzing results also need to be implemented separately by each user.^11, 12^ Lack of model standardization makes it hard to understand, reproduce and reuse many existing models and simulation results.

We introduce a new software tool, NetPyNE^†^. NetPyNE addresses these issues and relieves the user from much of the time-consuming coding previously needed for these ancillary modeling tasks, automating many network modeling requirements for the setup, run, explore and analysis stages. NetPyNE enables users to consolidate complex experimental data with prior models and other external data sources at different scales into a unified computational model. Users can then simulate and analyze the model in the NetPyNE framework in order to better understand brain structure, brain dynamics and ultimately brain structure-function relationships. The NetPyNE framework combines: **1.** flexible, rule-based, high-level standardized specifications covering scales from molecule to cell to network; **2.** efficient parallel simulation both on stand-alone computers and in high-performance computing (HPC) clusters; **3.** automated data analysis and visualization (*e.g.,* connectivity, neural activity, information theoretic analysis); **4.** standardized input/output formats, importing of existing NEURON cell models, and conversion to/from NeuroML;^13, 14^ **5.** automated parameter tuning (molecular to network levels) using grid search and evolutionary algorithms. All tool features are available programmatically or via an integrated graphical user interface (GUI). This centralized organization gives the user the ability to interact readily with the various components (for building, simulating, optimizing and analyzing networks), without requiring additional installation, setup, training and format conversion across multiple tools.

NetPyNE’s high-level specifications are implemented as a declarative language designed to facilitate the definition of data-driven multiscale network models by accommodating many of the intricacies of experimental data, such as complex subcellular mechanisms, the distribution of synapses across fully-detailed dendrites, and time-varying stimulation. Contrasting with the obscurity of raw-code descriptions used in many existing models,^15^ NetPyNE’s standardized language provides transparent and manageable descriptions. Model specifications are then translated into the necessary NEURON components via built-in algorithms. This approach cleanly separates model specifications from the underlying technical implementation. Users avoid complex low-level coding, preventing implementation errors, inefficiencies and flawed results that are common during the development of complex multiscale models. Crucially, users retain control of the model design choices, including the conceptual model, level of biological detail, scales to include, and biological parameter values. The NetPyNE tool allows users to shift their time, effort and focus from low-level coding to designing a model that matches the biological details at the chosen scales.

NetPyNE is one of several tools that facilitate network modeling with NEURON: neuroConstruct,^16^ PyNN,^17^ Topographica,^18^ ARACHNE^19^ and BioNet.^20^ NetPyNE differs from these in terms of the range of scales, from molecular up to large networks and extracellular space simulation – it is the only tool that supports NEURON’s Reaction-Diffusion (RxD) module.^21, 22^ It also provides an easy declarative format for the definition of complex, experimentally-derived rules to distribute synapses across dendrites. NetPyNE is also unique in integrating a standardized declarative language, automated parameter optimization and a GUI designed to work across all these scales.

NetPyNE therefore streamlines the modeling workflow, consequently accelerating the iteration between modeling and experiment. By reducing programming challenges, our tool also makes multiscale modeling highly accessible to a wide range of users in the neuroscience community. NetPyNE is publicly available from www.netpyne.org, which includes installation instructions, documentation, tutorials, example models and Q&A forums. The tool has already been used by over 40 researchers in different labs to train students and to model a variety of brain regions and phenomena (see www.netpyne.org/models).^23–26^ Additionally, it has also been integrated with other tools in the neuroscience community: the Human Neocortical Neurosolver (https://hnn.brown.edu/),^27^, ^28^ Open Source Brain^29^ (www.opensourcebrain.org),^14^ and the Neuroscience Gateway^30^ (www.nsgportal.org).

## 2 Results

### 2.1 Tool overview and workflow

NetPyNE’s workflow consists of four main stages: **1.** high-level specification, **2.** network instantiation, **3.** simulation and **4.** analysis and saving (Fig. 1). The first stage involves defining all the parameters required to build the network, from population sizes to cell properties to connectivity rules, and the simulation options, including duration, integration step, variables to record, *etc*. This is the main step requiring input from the user, who can provide these inputs either programmatically with NetPyNE’s declarative language, or by using the GUI. NetPyNE also enables importing of existing cell models for use in a network.

**Figure 1:**
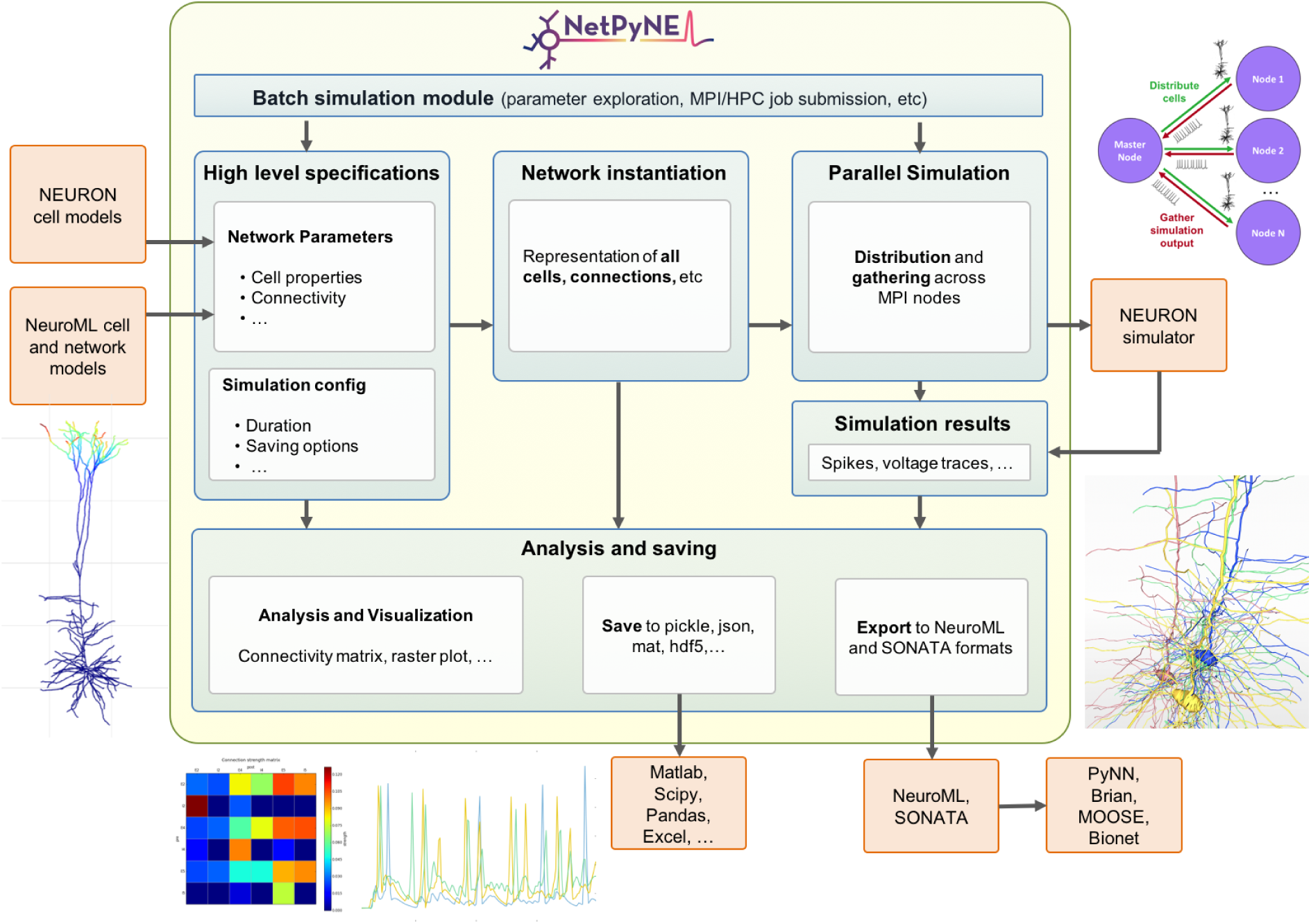
Overview of NetPyNE components and workflow. Users start by specifying the network parameters and simulation configuration using a high-level JSON-like format. Existing NEURON and NeuroML models can be imported. Next, a NEURON network model is instantiated based on these specifications. This model can be simulated in parallel using NEURON as the underlying simulation engine. Simulation results are gathered in the master node. Finally, the user can analyze the network and simulation results using a variety of plots; save to multiple formats or export to NeuroML. The Batch Simulation module enables automating this process to run multiple simulations on HPCs and explore a range of parameter values.

The next stages can be accomplished with a single function call – or mouse click if using the GUI. The network instantiation step consists of creating all the cells, connections and stimuli based on the high-level parameters and rules provided by the user. The instantiated network is represented as a Python hierarchical structure that includes all the NEURON objects required to run a parallel simulation. This is followed by the simulation stage, where NetPyNE takes care of distributing the cells and connections across the available nodes, running the parallelized simulation, and gathering the data back in the master node. Here, NetPyNE is using NEURON as its back-end simulator, but all the technical complexities of parallel NEURON are hidden to the user. In the final stage, the user can plot a wide variety of figures to analyze the network and simulation output. The model and simulation output can be saved to common file formats and exported to NeuroML, a standard description for neural models.^14^ This enables exploring the data using other tools (e.g. MATLAB) or importing and running the model using other simulators (*e.g.,* NEST).

An additional overarching component enables users to automate these steps to run batches of simulations to explore model parameters. The user can define the range of values to explore for each parameter and customize one of the pre-defined configuration templates to automatically submit all the simulation jobs on multi-processor machines or supercomputers.

Each of these stages is implemented in modular fashion to make it possible to follow different workflows such as saving an instantiated network and then loading and running simulations at a later time. The following sections provide additional details about each simulation stage.

### 2.2 High-level specifications

A major challenge in building models is combining the data from many scales. In this respect, NetPyNE offers a substantial advantage by employing a human-readable, clean, rule-based shareable declarative language to specify networks and simulation configuration. These standardized high-level specifications employ a compact JSON-compatible format consisting of Python lists and dictionaries (Fig. 2). The objective of the high-level declarative language is to allow users to accurately describe the particulars and patterns observed at each biological scale, while hiding all the complex technical aspects required to implement them in NEURON. For example, one can define a probabilistic connectivity rule between two populations, instead of creating potentially millions of cell-to-cell connections with Python or hoc for loops. The high-level language enables structured specification of all the model parameters: populations, cell properties, connectivity, input stimulation and simulation configuration.

**Figure 2:**
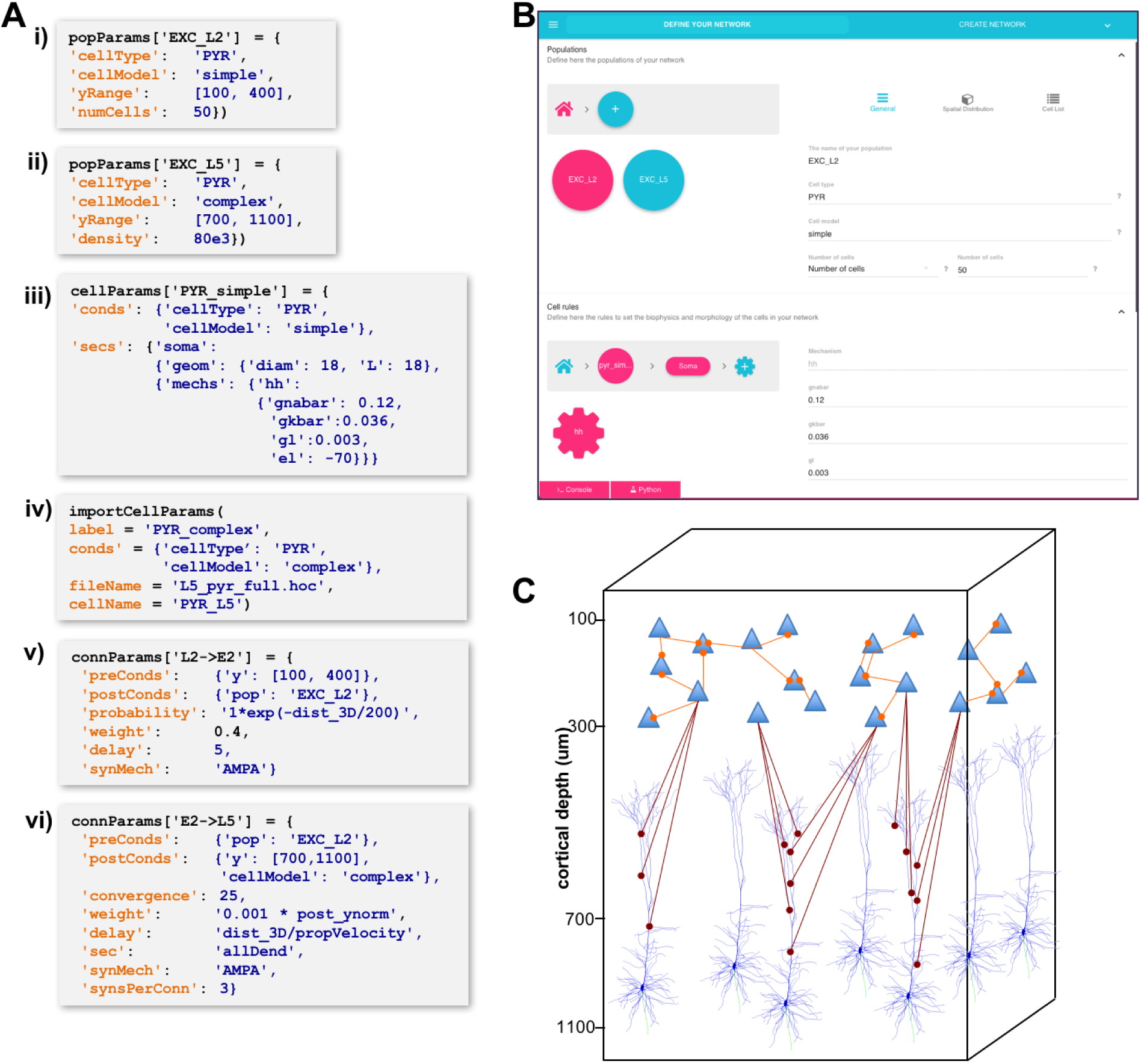
High-level specification of network parameters. A) Programmatic parameter specification using standardized declarative JSON-like format. i,ii: specification of two populations iii,iv: cell parameter definitions; v,vi connectivity rules. B) GUI-based parameter specification, showing the definition of populations equivalent to those in panel A. C) Schematic of network model resulting from the specifications in A.

#### 2.2.1 Population and cell parameters

Users define network populations, including their cell type, number of cells or density (in *cells/mm*^3^), and their spatial distribution. Fig. 2A-i,ii show setting of *yrange* and alternatively setting *numCells* or *density* for two cell types in the network. Morphological and biophysical properties can then be applied to subsets of cells using custom rules. This enables, for example, setting properties for all cells in a population with a certain “cell type” attribute or within a spatial region. The flexibility of the declarative rule-based method allows the heterogeneity of cell populations observed experimentally to be captured. It also allows the use of cell implementations of different complexity to coexist in the same network, useful in very large models where full multi-scale is desired but cannot be implemented across all cells due to the computational size of the network. These alternative implementations could include highly simplified cell models such as Izhikevich, AdEx or precalculated point neuron models.^31–33^ These can be combined in the same network model or swapped in and out: *e.g.,* **1.** explore overall network dynamics using simple point-neuron models; **2.** re-explore with more biologically realistic complex models to determine how complex cell dynamics contribute alters network dynamics. We also note that order of declaration is arbitrary; as here, one can define the density of typed cells before defining these types. In Fig. 2A-iii,iv, we define the two different *PY R* models whose distribution was defined in A-i,ii. The *simple* model is simple enough to be fully defined in NetPyNE – 1 compartment with Hodgkin-Huxley (*hh*) kinetics with the parameters listed (here the original *hh* parameters are given; typically these would be changed). More complex cells could also be defined in NetPyNE in this same way. More commonly, complex cells would be imported from hoc templates, Python classes or NeuroML templates, as shown in Fig. 2A-iv. Thus, any cell model available online can be downloaded and used as part of a network model (non-NEURON cell models must first be translated into NMODL/Python).^34^ Note that unlike the other statements, Fig. 2A-iv is a procedure call rather than the setting of a dictionary value. The `importCellParams()` procedure call creates a new dictionary with NetPyNE’s data structure, which can then be modified later in the script or via GUI, before network instantiation.

NetPyNE’s declarative language also supports NEURON’s reaction-diffusion RxD specifications of Regions, Species and Reactions.^21, 22^ RxD simplifies the declaration of the chemophysiology – intracellular and extracellular signaling dynamics – that complements electrophysiology. During network instantiation, RxD declarative specifications are translated into RxD components within or between cells of the NetPyNE-defined network. This adds additional scales – subcellular, organelle, extracellular matrix – to the exploration of multiscale interactions, *e.g.,* calcium regulation of HCN channels promoting persistent network activity.^35, 36^

#### 2.2.2 Connectivity and stimulation parameters

NetPyNE is designed to facilitate network design – connectivity rules are flexible and broad in order to permit ready translation of many different kinds of experimental observations. Different subsets of pre- and post-synaptic cells can be selected based on a combinations of attributes such as cell type and spatial location (Fig. 2A-v,vi) Users can then select the target synaptic mechanisms (*e.g.,* AMPA, AMPA/NMDA, GABA_A_). In the case of multicompartment cells, synapses can be distributed across a list of cell locations Multiple connectivity functions are available including all-to-all, probabilistic, fixed convergence and fixed divergence. The connectivity pattern can also be defined by the user via a custom connectivity matrix. Alternatively, connectivity parameters, typically including weight, probability and delay, can be specified as a function of pre- and post-synaptic properties. This permits instantiation of biological correlations such as the dependence of connection delay on distance, or a fall-off in connection probability with distance. Electrical gap junctions and learning mechanisms – including spike-timing dependent plasticity and reinforcement learning – can also be incorporated.

NetPyNE supports specification of subcellular synaptic distribution along dendrites. This allows synaptic density maps obtained via optogenetic techniques to be directly incorporated in networks. Fig. 3A left shows the layout for one such technique known as sCRACM (subcellular Channelrhodopsin-2-Assisted Circuit Mapping).^37^ A density map of cell activation measured from the soma is determined through light stimulation at the points on the grid in a slice whose presynaptic boutons from a particular projection, in this case from thalamus, have been tagged with channelrhodopsin (Fig. 3A). NetPyNE places synapses randomly based on location correspondence on a dendritic tree which can be either simple or multicompartmental (Fig. 3B). Here again, the automation of synapse placements permits models of different complexity to be readily swapped in and out. Depending on the data type and whether one wants to use averaging, the location maps may be based on 1D, 2D, or 3D tissue coordinates, with the major *y*-axis reflecting normalized cortical depth (NCD) from pia to white matter. Alternatively, NetPyNE can define synapse distributions based on categorical information for dendritic subsets: *e.g.,* obliques or spine densities, or on path distance from the soma, apical nexus or other point. As with the density maps, these rules will automatically adapt to simplified morphologies. NetPyNE permits visualization of these various synaptic-distribution choices and cellular models via dendrite-based synapse density plots (Fig. 3C), which in this case extrapolates from the experimental spatial-based density plot in Fig. 3A.^37–40^

**Figure 3:**
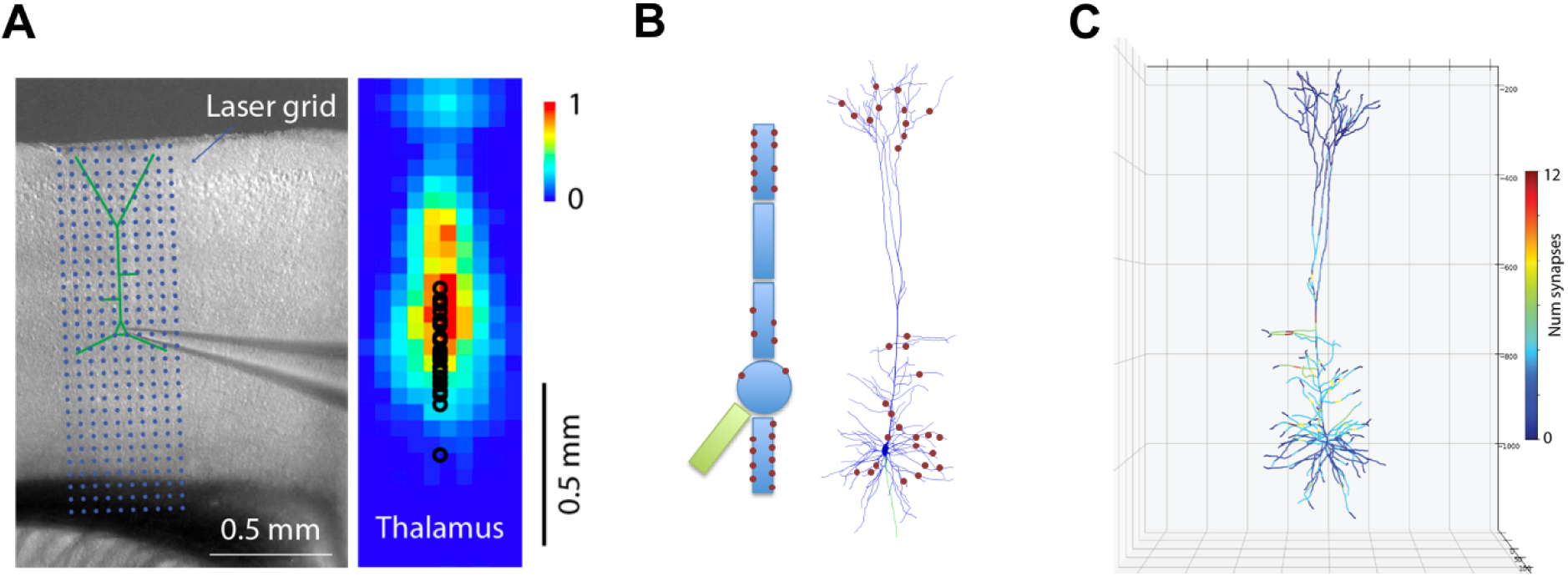
Specification of dendritic distribution of synapses. A) Optogenetic data provides synapse density across the 2D grid shown at left.^38^ B) Data are imported directly into NetPyNE which automatically calculates synapse location in simplified or full multicompartmental representations of a pyramidal cell. C) Corresponding synaptic density plot generated by NetPyNE.

Network models often employ artificial stimulation to reproduce the effect of afferent inputs that are not explicitly modeled, *e.g.,* ascending inputs from thalamus and descending from V2 targeting a V1 network. NetPyNE supports a variety of stimulation sources, including current clamps, random currents, random spike generators or band-delimited spike or current generators. These can be placed on target cells using the same flexible, customizable rules previously described for connections. Users can also employ experimentally recorded input patterns.

#### 2.2.3 Simulation configuration

Up to here, we have described the data structures, that defines network parameters: popParams, cellParams, connParams, *etc*. Next, the user will configure parameters related to a particular simulation run, such as simulation duration, time-step, parallelization options, *etc*. These parameters will also control output: variables to plot or to record for graphing – *e.g.,* voltage or calcium concentration from particular cells, LFP recording options, file save options, and in what format, *etc*. In contrast to network and cell parameterization, all simulation options have default values so only those being customized are required.

### 2.3 Network instantiation

NetPyNE generates a simulatable NEURON model containing all the elements and properties described by the user in the rule-based high-level specifications. As described above, declarations may include molecular processes, cells, connections, stimulators and simulation options. After instatiation, the data structures of both the original high-level specifications and the resultant network instance can be accessed programmatically or via GUI.

Traditionally, it has been up to the user to provide an easy way to access the components of a NEURON network model, *e.g.,* the connections or stimulators targeting a cell, the sections in a cell, or the properties and mechanisms in each section. This feature is absent in many existing models. Hence, inspecting these models requires calling multiple NEURON functions (*e.g.,* `SectionList.allroots(), SectionList.wholetree()` and `section.psection()`). Other models include some form of indexing for the elements at some scales, but, given this is not enforced, their structure and naming can vary significantly across models.

In contrast, all networks generated by NetPyNE are consistently represented as a nested Python structure. The root of the instantiated network is the *net* object (Fig. 4). *net* contains a list of cells; each cell contains lists or dictionaries with its properties, its sections, its stimulators. Each section *sec* contains dictionaries with its morphology and mechanisms. For example, once the network is instantiated, the sodium conductance parameter for cell#5 can be accessed as `net.cells[5].secs.soma.mechs.hh.gbar`. This data structure also includes all the NEURON objects – Sections, NetCons, NetStims, IClamps, *etc*. embedded hierarchically, and accessible via the `hObj` dictionary key of each element.

**Figure 4:**
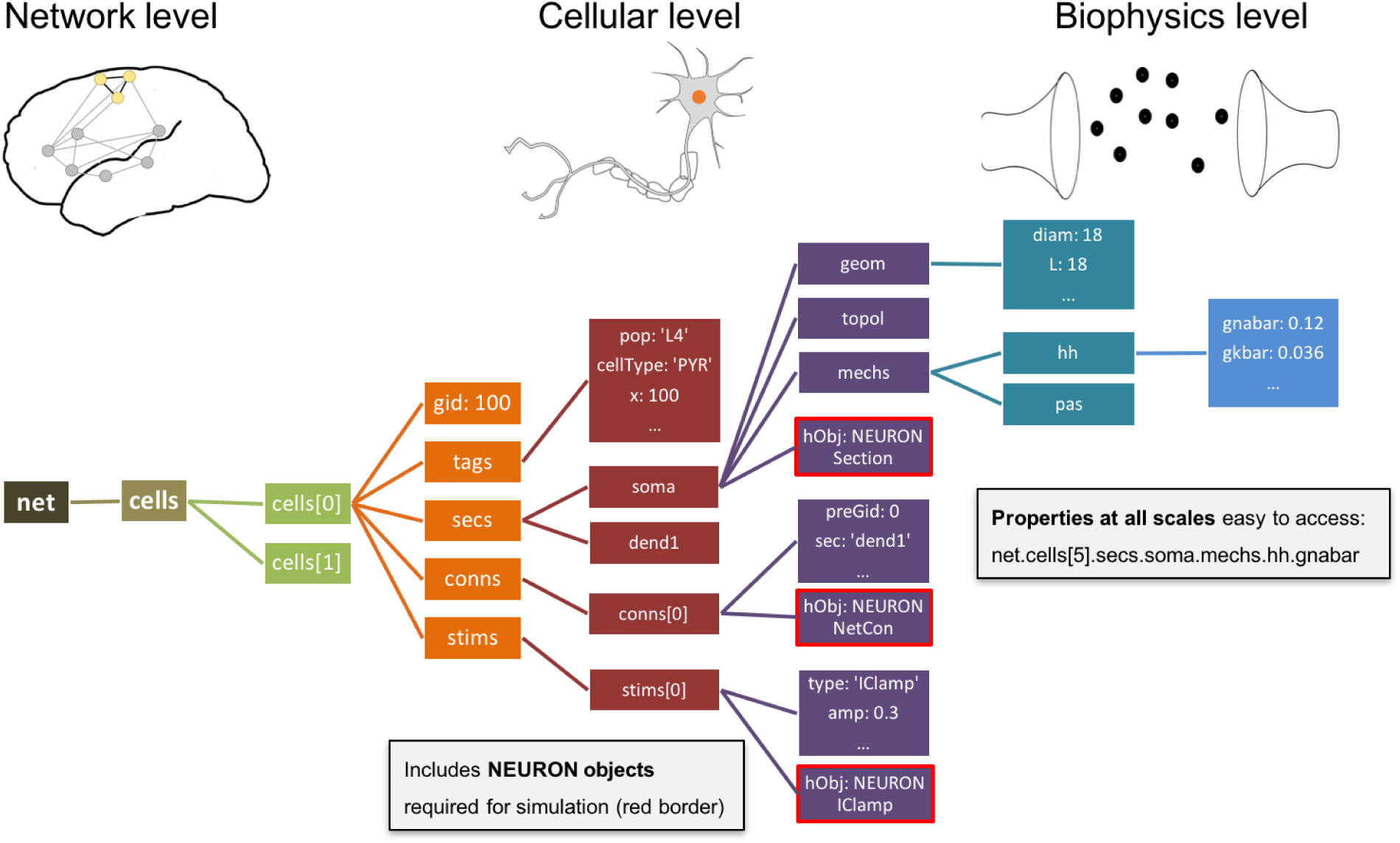
Instantiated network hierarchical data model. The instantiated network is represented using a standardized hierarchically-organized Python object that differs from the NetPyNE data structure of Fig. 1. Defined NEURON simulator objects are represented as boxes with red borders and correspond to the object type accessed via `h.objName`. These objects provide direct to all elements, state variables and parameters to be simulated.

### 2.4 Parallel simulation

Computational needs for running much larger and more complex neural simulations are constantly increasing as researchers attempt to reproduce fast-growing experimental datasets.^2, 4, 10, 23, 41, 42^ Fortunately, parallelization methods and high performance computing (HPC, supercomputing) resources are becoming increasingly available to the average user.^30, 43–48^

The NEURON simulator provides a *ParallelContext* module, which enables parallelizing the simulation computations across different nodes. However, this remains a complex process that involves distributing computations across nodes in a balanced manner, gathering and reassembling simulation results for post-processing, and ensuring simulation results are replicable and independent of the number of processors used. Therefore, appropriate and efficient parallelization of network simulations requires design, implementation and deployment of a variety of techniques, some complex, many obscure, mostly inaccessible to the average user.^10^

NetPyNE manages these burdensome tasks so that the user can run take a serial to a parallelized simulations with a single function call or mouse click. Cells are distributed across processors using a round-robin algorithm, which generally results in balanced computation load on each processor.^10, 49^ After the simulation has run, NetPyNE gathers in the master all the network metadata (cells, connections, *etc*.) and simulation results (spike times, voltage traces, LFP signal, *etc*.) for analysis. As models scale up, it becomes unfeasible to store the simulation results on a single centralized master node. NetPyNE offers distributed data saving methods that reduce both the runtime memory required and the gathering time. Distributed data saving means multiple compute nodes can write information in parallel, either at intervals during simulation runtime, or once the simulation is completed. The output files are later merged for analysis.

Random number generators (RNGs) are often problematic in hand-written parallelized code; careful management of seeds is required since use of the same seed or seed-sets across nodes will result in different random streams when the number of nodes is changed. Since random values are used to generate cell locations, connectivity properties, spike times of driving inputs, *etc*., inconsistent streams will cause a simulation to produce different results when going from serial to parallel or when changing the number of nodes. In NetPyNE, RNGs are initialized based on seed values created from associated pre- and post-synaptic cell global identifiers (gids) which ensures simulation stable results across different numbers of cores. Specific RNG streams are associated to *purposive* seeds (*e.g.,* connectivity or locations) and to a global seed, allowing different random, but replicable, networks to be run with change of the single global seed. Similarly, manipulation of *purposive* seeds can be used to run, for example, a network with identical wiring but different random driving inputs.

We previously performed parallelization performance analyses that demonstrated run time scales appropriately as a function of number of cells (tested up to 100,000) and compute nodes (tested up to 512).^10^ Simulations were developed and executed using NetPyNE and NEURON on the XSEDE Comet supercomputer via the Neuroscience Gateway^30^ (www.nsgportal.org). The Neuroscience Gateway, which provides neuroscientists with free and easy access to supercomputers, includes NetPyNE as one of the tools available via their web portal. Larger-scale models – including the M1 model with 10k multicompartment neurons and 30 million synapses^23^ and the thalamocortical model with over 80k point neurons and 300 million synapses^24, 50^ – have been simulated in both the XSEDE Comet supercomputer and Google Cloud supercomputers. Run time to simulate one second of the multicompartment-neuron network required 47 minutes on 48 cores, and 4 minutes on 128 cores for the point-neuron network.

### 2.5 Analysis of network and simulation output

To extract conclusions from neural simulations it is necessary to use further tools to process and present the large amounts of raw data generated. NetPyNE includes built-in implementations of a wide range of visualization and analysis functions commonly used in neuroscience (Fig. 5). All analysis functions include options to customize the desired output. Functions to visualize and analyze network structure are available without a simulation run: **1.** intracellular and extracellular RxD species concentration in a 2D region; **2.** matrix or stacked bar plot of connectivity; **3.** 2D representation of cell locations and connections; and **4.** 3D cell morphology with color-coded variable (*e.g.,* number of synapses per segment). After a simulation run, one can visualize and analyze simulation output: **1.** time-resolved traces of any recorded cell variable (*e.g.,* voltage, synaptic current or ion concentration); **2.** relative and absolute amplitudes of post-synaptic potentials; **3.** spiking statistics (boxplot) of rate, the interspike interval coefficient of variation (ISI CV) and synchrony;^51^ **4.** power spectral density of firing rates; and **5.** information theory measures, including normalized transfer entropy and Granger causality.

**Figure 5:**
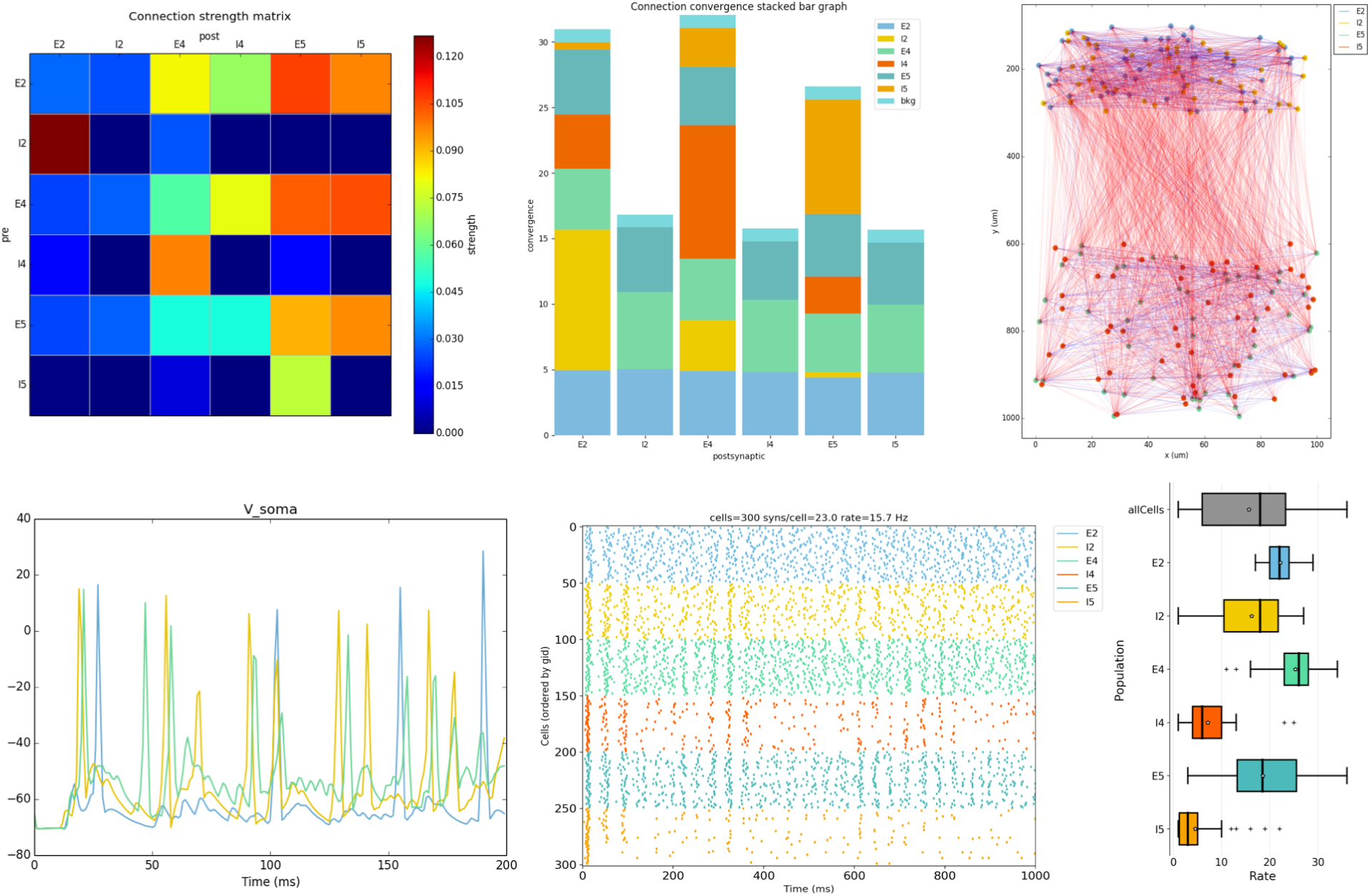
NetPyNE visualization and analysis plots for a simple 3-layer network example. A) Connectivity matrix, B) stacked bar graph, C) 2D representation of cells and connnetions, D) voltage traces of 3 cells, E) raster plot, F) population firing rate statistics (boxplot).

A major feature of our tool is the ability to place extracellular electrodes to record LFPs at any arbitrary 3D locations within the network, similar to the approach offered by the LFPy^52^ and LFPsim^53^ add-ons to NEURON. The LFP signal at each electrode is obtained by summing the extracellular potential contributed by each neuronal segment, calculated using the “line source approximation” and assuming an Ohmic medium with conductivity.^53, 54^ The user can then plot the location of each electrode, together with the recorded LFP signal and its power spectral density and spectrogram (Fig. 6). The ability to record and analyze LFPs facilitates reproducing experimental datasets that include this commonly used measure.^54^

**Figure 6:**
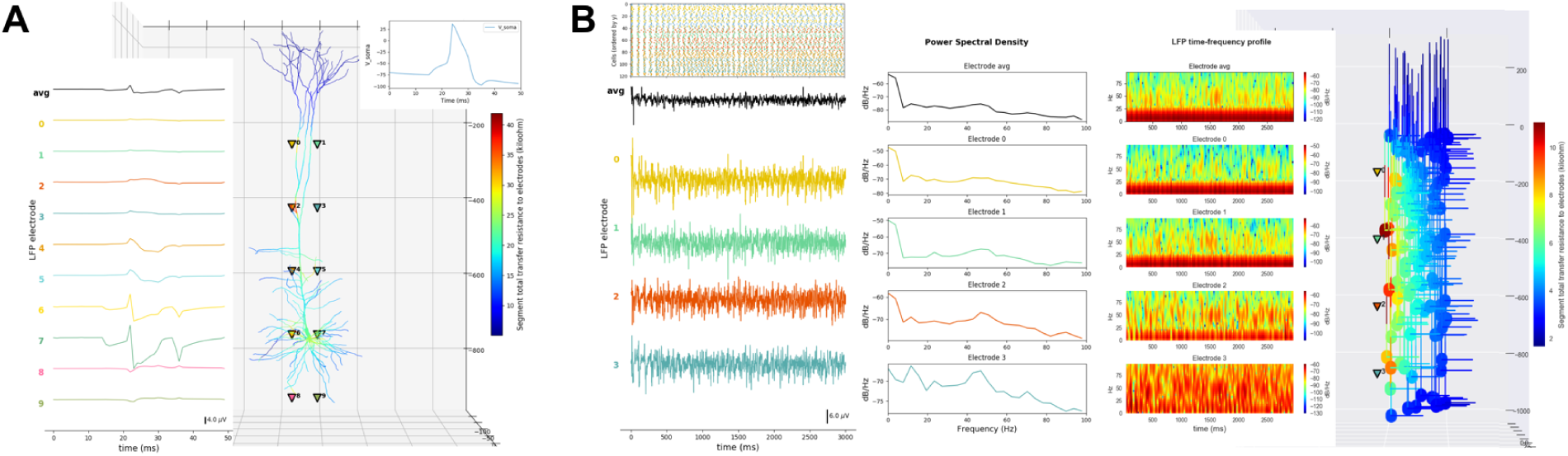
LFP recording and analysis. A) LFP signals (left) from 10 extracellular recording electrodes located around a morphologically detailed cell (right) producing a single action potential (top-right). B) LFP signals, PSDs and spectrograms (left and center) from 4 extracellular recording electrodes located at different depths of a network of 120 5-compartment neurons (right) producing oscillatory activity (top-left).

### 2.6 Data saving and exporting

NetPyNE permits saving and loading of all model components and results separately or in combination: high-level specifications, network instance, simulation configuration, simulation data, and simulation analysis results. Saving network instances enables loading a specific saved network with all explicit cells and connections, without the need to re-generate these from the high-level connectivity rules. NetPyNE supports several standard file formats: pickle, JSON, MAT, and HDF5. The use of common file formats allows network structure and simulation results to be easily analyzed using other tools such as MATLAB or Python Pandas.

Network instances can also be exported to or imported from NeuroML,^14^ a standard declarative format for neural models, and SONATA (https://github.com/AllenInstitute/sonata), a format standard for neural models proposed by the Blue Brain Project and Allen Institute for Brain Science. These formats are also supported by other simulation tools, so that models developed using NetPyNE can be exported, explored and simulated in other tools including Brian,^55^ MOOSE,^56, 57^ PyNN,^17^ Bionet^20^ or Open Source Brain.^29^ Similarly simulations from these other tools can be imported into NetPyNE. This feature also enables any NetPyNE model to be visualized via the Open Source Brain portal, and permits a NeuroML model hosted on the portal to be parallelized across multiple cores (*e.g.,* on HPC) using NetPyNE.

Long simulations of large networks take a long time to run. Due to memory and disk constraints, it is not practical to save all state variables from all cells during a run, particularly when including signaling concentrations at many locations when using the reaction-diffusion module. Therefore, NetPyNE includes the option of recreating single cell activity in the context of spike inputs previously recorded from a network run. These follow-up simulations do not typically require an HPC since they are only running the one cell. The user selects a time period, a cell number, and a set of state variables to record or graph.

### 2.7 Parameter optimization and exploration via batch simulations

Parameter optimization involves finding sets of parameters that lead to a desired output in a model. This process is often required since both single neuron and network models include many not-fully constrained parameters that can be modified within a known biological range of values. Network dynamics can be highly sensitive, with small parameter variations leading to large changes. This then requires searching within complex multidimensional spaces to match experimental data, with degeneracy such that multiple parameter sets may produce matching activity patterns.^58–60^ A related concept is that of parameter exploration. Once a model is tuned to reproduce biological features, it is common to explore individual parameters to understand their relation to particular model features, *e.g.,* how synaptic weights affect network oscillations,^61^ or the effect of different pharmacological treatments on pathological symptoms.^26, 62^

Many different approaches exist to perform parameter optimization and exploration. Manual tuning usually requires expertise and a great deal of patience.^63, 64^ Therefore, NetPyNE provides built-in support for several automated methods that have been successfully applied to both single cell and network optimization: grid-search and various types of evolutionary algorithms (EAs).^2, 65–70^ Grid search refers to evaluating combinations on a fixed set of values for a chosen set of parameters, resulting in gridded sampling of the multidimensional parameter space. EAs search parameter space more widely and are computationally efficient when handling complex, non-smooth, high-dimensional parameter spaces.^64^ They effectively follow the principles of biological evolution: here a population of models evolves by changing parameters in a way that emulates crossover events and mutation over generations until individuals reach a desired fitness level.

NetPyNE provides an automated parameter optimization and exploration framework specifically tailored to multiscale biophysically-detailed models. Our tool facilitates the multiple steps required: **1.** parameterizing the model and selecting appropriate value ranges; **2.** providing a fitness functions; **3.** customizing the optimization/exploration algorithm options; **4.** running the batch simulations; and **5.** managing and analyzing batch simulation parameters and outputs. To facilitate parameter selection, all of the network specifications are available to the user via the NetPyNE declarative data structure – from molecular concentrations and ionic channel conductances to long-range input firing rates – freeing the user from having to identify parameters or state variables at the simulator level.

Both parameter optimization and exploration involve running many instances of the network with different parameter values, and thus typically require parallelization. For these purposes, NetPyNE parallelization is implemented at two levels: **1.** simulation level – cell computations distributed across nodes as described above; and **2.** batch level – many simulations with different parameters executed in parallel.^65^ NetPyNE includes predefined execution setups to automatically run parallelized batch simulations on different environments: **1.** multiprocessor local machines or servers via standard message passing interface (MPI) support; **2.** the Neuroscience Gateway (NSG) online portal, which includes compressing the files and uploading a zip file via RESTful services; **3.** HPC systems (supercomputers) that employ job queuing systems such as PBS Torque or SLURM (*e.g.,* Google Cloud Computing HPCs). Users will be able to select the most suitable environment setup and customize options if necessary, including any optimization algorithm metaparameters such as population size, mutation rate for EAs. A single high-level command will then take care of launching the batch simulations to optimize or to explore the model.

### 2.8 Graphical User Interface (GUI)

The GUI enables users to more intuitively access NetPyNE functionalities. It divides the workflow into two tabs: network definition and network exploration, simulation and analysis. From the first tab it is possible to define – or import from various formats – the high-level network parameters/rules and simulation configuration (Fig. 2B). Parameter specification is greatly facilitated by having clearly structured and labeled sets of parameters, graphics to represent different components, drop-down lists, autocomplete forms and automated suggestions. The GUI also includes an interactive Python console and full bidirectional synchronization with the underlying Python-based model – parameters changed via the Python console will be reflected in the GUI, and vice versa. In the second tab the user can interactively visualize the instantiated network in 3D, run parallel simulations and display all the available plots to analyze the network and simulation results. An example of a multiscale model visualized, simulated and analyzed using the GUI is shown in Fig. 7. The code and further details of this example are available at https://github.com/Neurosim-lab/netpyne/tree/development/examples/rxd_net.

**Figure 7:**
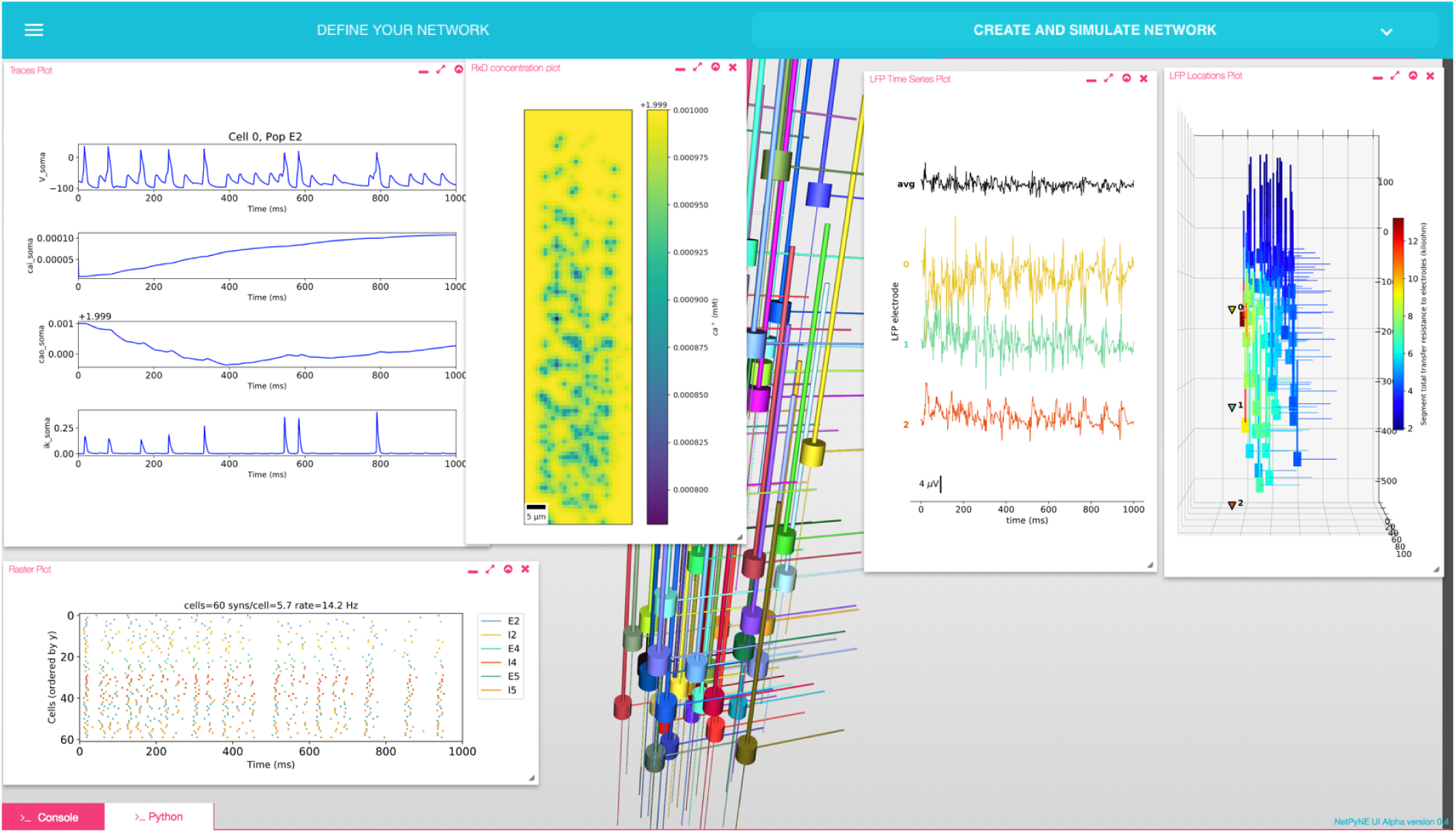
NetPyNE graphical user interface (GUI) showing a multiscale model. Background shows 3D representation of example network with 6 populations of multi-channel multi-compartment neurons (back-ground); plots from left to right: cell traces (voltage, intracellular and extracellular calcium concentration, and potassium current); raster plot; extracellular potassium concentration; LFP signals recorded from 3 electrodes; and 3D location of the LFP electrodes within network.

The GUI is particularly useful for beginners, students or non-computational researchers who can rapidly build networks without knowledge of coding and without learning NetPyNE’s declarative syntax. From there, they can simulate and explore multiscale subcellular, cellular and network models with varying degrees of complexity, from integrate-and-fire up to large-scale simulations that require HPCs. The GUI is also useful for modelers, who can easily prototype new models graphically and later extend the model programmatically using automatically generated Python scripts. Finally, the GUI is useful – independently of expertise level – to explore and visualize existing models developed by oneself, developed by other users programmatically, or imported from other simulators. Understanding unfamiliar models is easier if users can navigate through all the high-level parameters in a structured manner and visualize the instantiated network structure, instead of just looking at the model definition code.^71^

### 2.9 Application examples

Our recent model of primary motor cortex (M1) microcircuits^23, 26, 66^ constitutes an illustrative example where NetPyNE enabled the integration of complex experimental data at multiple scales: it simulates over 10,000 biophysically detailed neurons and 30 million synaptic connections. Neuron densities, classes, morphology and biophysics, and connectivity at the long-range, local and dendritic scale were derived from published experimental data.^38–40, 72, 73, 73–79^ Results yielded insights into circuit information pathways, oscillatory coding mechanisms and the role of HCN in modulating corticospinal output.^23^ A scaled down version (180 neurons) of the M1 model is illustrated Fig. 8.

**Figure 8:**
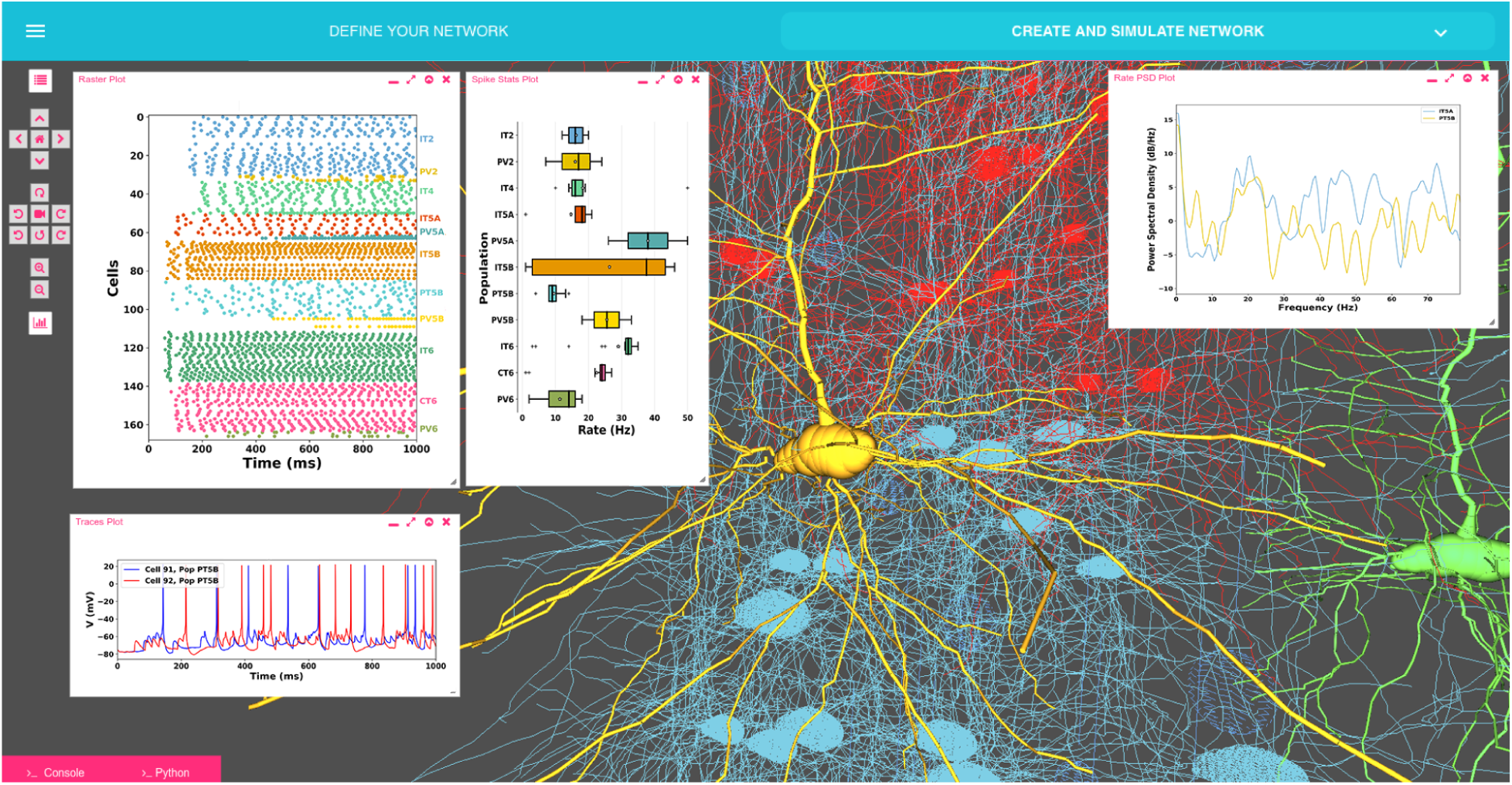
Model of M1 microcircuits developed using NetPyNE (scaled down version). NetPyNE GUI showing 3D representation of M1 network (background), raster plot and population firing rate statistics (top left), voltage traces (bottom left) and firing rate power spectral density (top right).

Several models published in other languages have been converted into NetPyNE to increase their usability and flexibility. These include models of cortical circuits exploring EEG/MEG signals (https://hnn.brown.edu/),^27^, ^28^ interlaminar flow of activity^24, 50^ (Fig. 9A) and epileptic activity^62^ (Fig. 9B); a dentate gyrus network^80, 81^ (Fig. 9C); and CA1 microcircuits^82, 83^ (Fig. 9D). As a measure of how compact the model definition is, we compared the number of source code lines (excluding comments, blank lines, cell template files and mod files) of the original and NetPyNE implementations (see Table 2.9).

**Figure 9:**
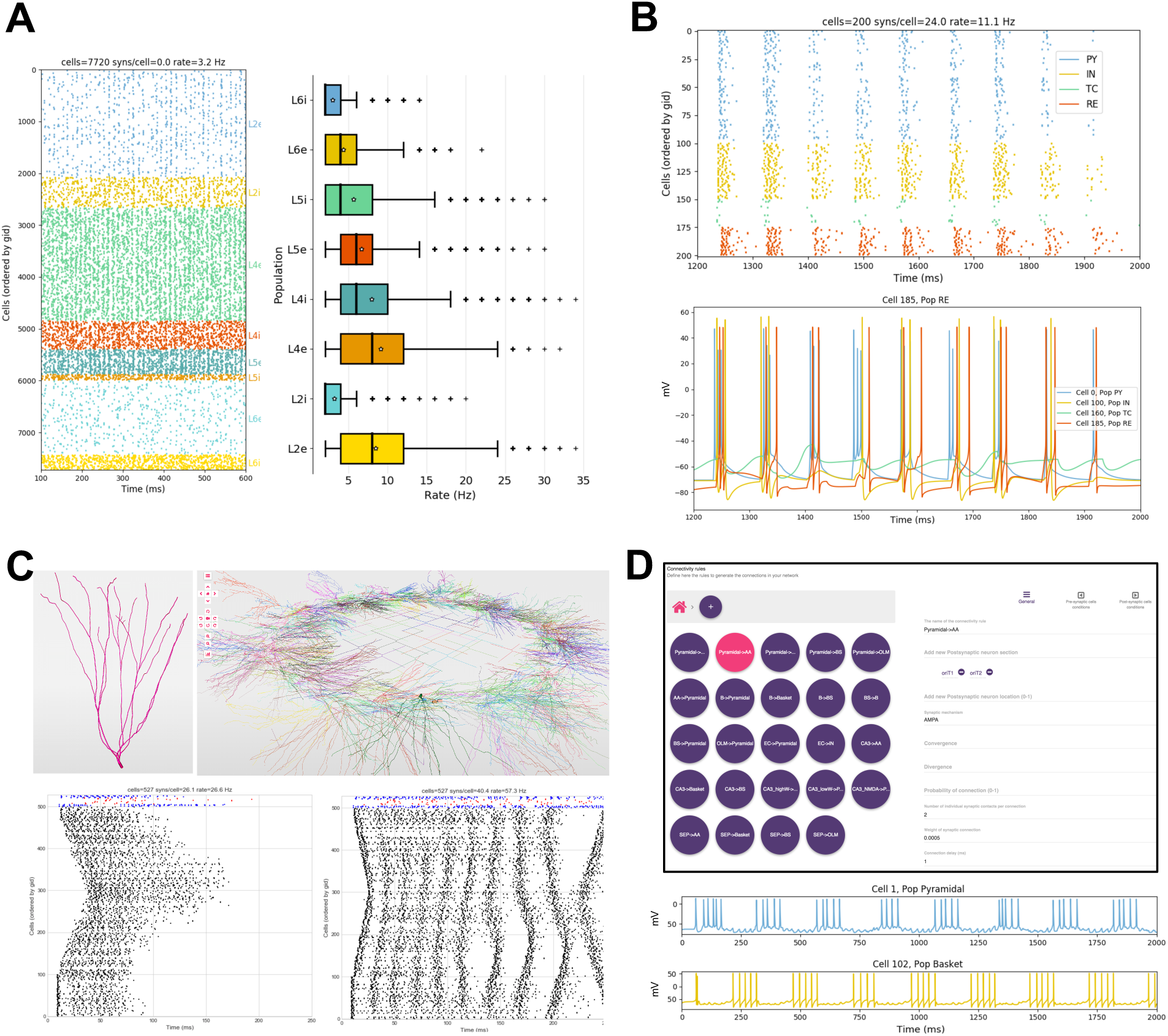
Published models converted to NetPyNE. All figures were generated using the NetPyNE version of the models. A) Raster plot and boxplot statistics of the Potjans and Diesmann thalamocortical network originally implemented in NEST.^24, 50^ B) Raster plot and voltage traces of a thalamocortical network exhibiting epileptic activity originally implemented in NEURON/hoc.^62^ C) 3D representation of the cell types and network topology, and raster plots of dentate gyrus model originally implemented in NEURON/hoc.^80, 81^ D) Connectivity rules (top) and voltage traces of 2 cell types (bottom) of a hippocampal CA1 model originally implemented in NEURON/hoc.^82, 83^

| Model description | Original language | Original num lines | NetPyNE num lines |
| --- | --- | --- | --- |
| Dentate gyrus<sup>80</sup> | NEURON/hoc | 1029 | 261 |
| CA1 microcircuits<sup>82</sup> | NEURON/hoc | 642 | 306 |
| Epilepsy in thalamocortex<sup>62</sup> | NEURON/hoc | 556 | 201 |
| EEG/MEG in cortex (HNN model)<sup>27,28</sup> | NEURON/Python | 2288 | 924 |
| Motor cortex with RL<sup>65</sup> | NEURON/Python | 1171 | 362 |
| Cortical microcircuits<sup>50</sup> | PyNEST | 689 | 198 |

## 3 Discussion

NetPyNE is a high-level Python interface to the NEURON simulator that facilitates the definition, parallel simulation, optimization and analysis of data-driven brain circuit models. NetPyNE provides a systematic, standardized approach to biologically-detailed multiscale modeling. Its broad scope offers users the option to evaluate neural dynamics from a variety of scale perspectives: *e.g.,* **1.** network simulation in context of the brain as an organ – *i.e.,* with extracellular space included; **2.** focus at the cellular level in the context of the network; **3.** evaluate detailed spine and dendrite modeling in the context of the whole cell *and* the network, *etc*. Swapping focus back-and-forth across scales allows the investigator to understand scale integration in a way that cannot be done in the experimental preparation. In this way, multiscale modeling complements experimentation by combining and making interpretable previously incommensurable datasets. *In silico* models developed with NetPyNE can serve as testbeds that can be probed extensively and precisely in ways that parallel experimentation to make testable predictions. Simulation can also go beyond the capabilities of physical experimentation to build comprehension and develop novel theoretical constructs.^2, 4, 41, 84^

To ensure accessibility to a wide range of researchers, including modelers, students and experimentalists, NetPyNE combines many of the modeling workflow features under a single framework with both a programmatic and graphical interface. The GUI provides an intuitive way to learn to use the tool and explore all the different components and features interactively. Exporting the generated network to a Python script enables more advanced users to extend the model programmatically.

### 3.1 Multiscale specifications using a declarative language

By providing support for NEURON’s intracellular and extracellular reaction-diffusion module (RxD),^21, 22^ NetPyNE helps to couple molecular-level chemophysiology – historically neglected in computational neuroscience – to the classical electrophysiology at subcellular, cellular and network scales. RxD allows the user to specify and simulate the diffusion of molecules (*e.g.,* calcium, potassium or IP3) intracellularly, subcellularly (by including organelles such as endoplasmic reticulum and mitochondria), and extracellularly in the context of signaling and enzymatic processing – *e.g.,* metabolism, phosphorylation, buffering, second messenger cascades. This relates the scale of molecular interactions with that of cells and networks.

NetPyNE rules allow users to not only define connections at the cell-to-cell level, but also to compactly express highly specific patterns of the subcellular distribution of synapses, *e.g.,* depending on the neurite cortical depth or path distance from soma. Such distinct innervation patterns have been shown to depend on brain region, cell type and location and are likely to subserve important information processing functions and have effects at multiple scales.^37, 39, 85, 86^ Some simulation tools (GENESIS,^56^ MOOSE, PyNN^17^ and neuroConstruct^16^) include basic dendritic level connectivity features, and others (BioNet^20^) allow for Python functions that describe arbitrarily complex synapse distribution and connectivity rules. However, NetPyNE is unique in facilitating the description of these synaptic distribution patterns via flexible high-level declarations that require no algorithmic coding.

NetPyNE’s high-level language has advantages over procedural description in that it provides a human-readable, declarative format, accompanied by a parallel graphical representation, making models easier to read, modify, share and reuse. Other simulation tools such as PyNN, NEST, Brian or BioNet include high-level specifications in the context of the underlying procedural language used for all aspects of model instantiation, running and initial analysis. Procedural languages require ordering by the logic of execution rather than the logic of the conceptual model. Since the NetPyNE declarative format is order free, it can be cleanly organized by scale, by cell type, or by region at the discretion of the user. This declarative description can then be stored in standardized formats that can be readily translated into shareable data formats for use with other simulators. High-level specifications are translated into a network instance using previously tested and debugged implementations. Compared to creating these elements directly via procedural coding (in Python/NEURON), our approach reduces the chances of coding bugs, replicability issues and inefficiencies,

The trade-off is that users of a declarative language are constrained to express inputs according to the standardized formats provided, offering somewhat less flexibility compared to a procedural language. However, NetPyNE has been designed so that many fields are agglutinative, allowing multiple descriptors to be provided together to hone in on particular subsets of cells, subcells or subnetworks, *e.g.,* cells of a certain type within a given spatial region. Additionally, users can add procedural NEURON/Python code between the instantiation and simulation stages of NetPyNE in order to customize or add non-supported features to the model.

Developers of several applications and languages, including NeuroML, PyNN, SONATA and NetPyNE, are working together to ensure interoperability between their different formats. NeuroML^14^ is a widely-used model specification language for computational neuroscience which can store instantiated networks through an explicit list of populations of cells and their connections, without higher level specification rules. We are collaborating with the NeuroML developers to incorporate high-level specifications similar to those used in NetPyNE, *e.g.,* compact connectivity rules (see github.com/NeuroML/NeuroMLlite). The hope is that these compact network descriptions become a standard in the field so that they can be used to produce identical network instances across different simulators. To further promote standardization and interoperability, we and other groups working on large-scale networks founded the INCF Special Interest Group on “Standardized Representations of Network Structures” (www.incf.org/activities/standards-and-best-practices/incf-special-interest-groups/incf-sig-on-standardised).

### 3.2 Integrated parameter optimization

A major difficulty in building complex models is optimizing its many parameters within biological constrains to reproduce experimental results.^63, 64^ Multiple tools are available to fit detailed single cell models to electrophysiological data: BluePyOpt,^87^ Optimizer,^88^ Pypet^89^ or NeuroTune.^90^ However, these optimizers work within a single scale rather than optimizing across scales to study complex cells in complex circuits. NetPyNE provides a parameter optimization framework designed specifically to tackle this problem, thus enabling and encouraging the exploration of interactions across scales. It also closely integrates with the simulator rather than being a standalone optimizer, which would require expertise to interface properly. NetPyNE offers multiple optimization methods, including evolutionary algorithms, which are computationally efficient for handling the non-smooth high-dimensional parameter spaces found in this domain.^63, 64, 91^

### 3.3 Use of NetPyNE in education

In addition to the tool itself, we have developed detailed online documentation, step-by-step tutorials (www.netpyne.org), and example models. The code has been released as open source (github.com/Neurosim-lab/netpyne). Ongoing support is provided via a mailing list (with 50 subscribed users) and active Q&A forums (150 posts and over 5,000 views in the first year): www.netpyne.org/mailing, www.netpyne.org/forumandnetpyne.org/neuron-forum. Users have been able to quickly learn to build, simulate and explore models that illustrate fundamental neuroscience concepts, making NetPyNE a useful tool to train students. To disseminate the tool we have also provided NetPyNE training at conference workshops and tutorials, summer schools and university courses. Several labs are beginning to use NetPyNE to train students and postdocs.

### 3.4 Use of NetPyNE in research

Models being developed in NetPyNE cover a wide range of regions including thalamus, sensory and motor cortices,^23, 26^ claustrum,^25^ striatum, cerebellum, hippocampus. Application areas being explored include schizophrenia, epilepsy, transcranial magnetic stimulation (TMS), and electro- and magneto-encephalography (EEG/MEG) signals.^92^ A full list of areas and applications is available at www.netpyne.org/models.

Tools such as NetPyNE that provide insights into multiscale interactions are particularly important for the understanding of brain disorders, which always involve interactions across spatial and temporal scale domains.^93^ Development of novel biomarkers, increased segregation of disease subtypes, new treatments, and personalized treatments, all require that details of molecular, anatomical, functional, and dynamic organization that have been demonstrated in isolation be related to one another. Simulations and analyses developed in NetPyNE provide a way to link these scales, from the molecular processes of pharmacology, to cell biophysics, electrophysiology, neural dynamics, population oscillations, EEG/MEG signals and behavioral measures.

## 4 Methods

### 4.1 Overview of tool components and workflow

NetPyNE is implemented as a Python package that acts as a high-level interface to the NEURON simulator. The package is divided into several subpackages, which roughly match the components depicted in the workflow diagram in Fig. 1. The specs subpackage contains modules related to definition of high-level specifications. The sim subpackage contains modules related to running the simulation. It also serves as a shared container that encapsulates and provides easy access to the remaining subpackages, including methods to build the network or analyze the output, and the actual instantiated network and cell objects. From the user perspective, the basic modeling workflow is divided into three steps: defining the network parameters (populations, cell rules, connectivity rules, *etc*) inside an object of the class `specs.NetParams;` setting the simulation configuration options (run time, integration interval, recording option, *etc*) inside an object of the class `specs.SimConfig;` and passing these two objects to a wrapper function (`sim.createSimulateAnalyze()`) that takes care of creating the network, running the simulation and analyzing the output.

### 4.2 Network instantiation

The following standard sequence of events are executed internally to instantiate a network from the high-level specifications in the `netParams` object: **1.** create a `Network` object and add to it a set of `Population` and `Cell` objects based on parameters; **2.** set cell properties (morphology and biophysics) based on `cellParams` parameters (checking which cells match the conditions of each rule); **3.** create molecular-level RxD objects based on `rxdParams` parameters; **4.** add stimulation (IClamps, NetStims, *etc*) to the cells based on `stimSourceParams` and `stimTargetParams` parameters; and **5.** create a set of connections based on `connParams` and `subConnParams` parameters (checking which presynpatic and postsynaptic cells match the conn rule conditions), with the synaptic parameters specified in `synMechParams`. After this process is completed all the resulting NEURON objects will be contained and easily accessible within a hierarchical Python structure (object `sim.net` of the class `Network`) as depicted in Fig. 4.

The network building task is further complicated by the need to implement parallel NEURON simulations in an efficient and replicable manner, independent of the number of processors employed. Random number generators (RNGs) are used in several steps of the building process, including cell locations, connectivity properties and the spike times of input stimuli (*e.g.,* NetStims). To ensure random independent streams that can be replicated deterministically when running on different number of cores we employed NEURON’s Random123 RNG from the `h.Random` class. This versatile cryptographic quality RNG^94^ is initialized using three seed values, which, in our case, will include a global seed value and two other values related to unique properties of the cells involved, *e.g.,* for probabilistic connections, the gids of the pre- and post-synaptic cells.

To run NEURON parallel simulations NetPyNE employs a pc object of the class `h.ParallelContext()`, which is created when the sim object is first initialized. During the creation of the network, the cells are registered via the pc methods to enable exchange and recording of spikes across compute nodes. Prior to running the simulation, global variables, such as temperature or initial voltages are initialized, and the recording of any traces (*e.g.,* cell voltages) and LFP is set up by creating `h.Vector()` containers and calling the recording methods. After running the parallel simulation via `pc.solve()`, data (cells, connections, spike times, recorded traces, LFPs, *etc*) is gathered into the master node from all compute nodes using the `pc.py alltoall()` method. Alternatively, distributed saving enables writing the output of each node to file and combining these files after the simulation has ended. After gathering, the built-in analysis functions have direct access to all the network and simulation output data via `sim.net.allCells` and `sim.allSimData`.

### 4.3 Importing and exporting

NetPyNE enables importing existing cells in hoc or Python, including both templates/classes and instantiated cells. To do this NetPyNE internally runs the hoc or Python cell model, extracts all the relevant cell parameters (morphology, mechanisms, point processes, synapses, *etc*) and stores them in the NetPyNE JSON-like format used for high-level specifications. The hoc or Python cell model is then completely removed from memory so later simulations are not affected.

Importing and exporting to other formats such as NeuroML or SONATA requires mapping the different model components across formats. To ensure validity of the conversion we have compared simulation outputs from each tool, or converted back to the original format and compared to the original model. Tests on mappings between NetPyNE and NeuroML can be found at https://github.com/OpenSourceBrain/NetPyNEShowcase.

### 4.4 Batch simulations

Exploring or fitting model parameters typically involves running many simulations with small variations in some parameters. NetPyNE facilitates this process by automatically modifying these parameters and running all the simulations based on a set of high-level instructions provided by the user. The two fitting approaches – grid search and evolutionary algorithms – both require similar set up. The user creates a `Batch` object that specifies the range of parameters values to be explored and the run configuration (*e.g.,* use 48 cores on a cluster with SLURM workload manager). For evolutionary algorithms and optionally for grid search, the user provides a Python function that acts as the algorithm fitness function, which can include variables from the network and simulation output data (*e.g.,* average firing rate of a population). The tool website includes documentation and examples on how to run the different types of batch simulations.

Once the batch configuration is completed, the user can call the `Batch.run()` method to trigger the execution of the batch simulations. Internally, NetPyNE iterates over the different parameter combinations. For each one, NetPyNE will **1.** set the varying parameters in the simulation configuration (`SimConfig` object) and save it to file, **2.** launch a job to run the NEURON simulation based on the run options provided by the user (*e.g.,* submit a SLURM job), **3.** store the simulation output with a unique filename, and **4.** repeat for the next parameter set, or if using evolutionary algorithms, calculate the fitness values and the next generation of individuals (parameter sets).

To implement the evolutionary algorithm optimization we made use of the Inspyred Python package (https://pythonhosted.org/inspyred/). Inspyred subroutines are particularized to the neural environment, directly using parameters and fitness values obtained from NetPyNE data structures, and running parallel simulations under the NEURON environment either in multiprocessor machines via MPI or supercomputers via workload managers.

### 4.5 Graphical User Interface

The NetPyNE GUI is implemented on top of Geppetto,^95^ an open-source platform that provides the infrastructure for building tools for visualizing neuroscience models and data and managing simulations in a highly accessible way. The GUI is defined using Javascript, React and HTML5. This offers a flexible and intuitive way to create advanced layouts while still enabling each of the elements of the interface to be synchronized with the Python model. The interactive Python backend is implemented as a Jupyter Notebook extension which provides direct communication with the Python kernel. This makes it possible to synchronize the data model underlying the GUI with a custom Python-based NetPyNE model. This functionality is at the heart of the GUI and means any change made to the NetPyNE model in Python kernel is immediately reflected in the GUI and vice versa. The tool’s GUI is available at https://github.com/MetaCell/NetPyNE-UI and is under active development.

## Acknowledgements

This work was funded by the following grants: NIH grant U01EB017695, DOH01-C32250GG-3450000, NIH R01EB022903, NIH R01MH086638, and NIH 2R01DC012947-06A1. PG was funded by the Wellcome Trust (101445). We are thankful to all the contributors that have collaborated in the development of this open source tool via GitHub (https://github.com/Neurosim-lab/netpyne).

## Footnotes

† NetPyNE: Network specification, simulation and analysis using Python and NEURON.

## References

1. Cornelia Bargmann, William Newsome, A Anderson, E Brown, K Deisseroth, J Donoghue, P MacLeish, E Marder, R Normann, J Sanes, et al. Brain 2025: a scientific vision. Brain Research through Advancing Innovative Neurotechnologies (BRAIN) Working Group Report to the Advisory Committee to the Director, NIH, 2014.

2. Henry Markram, Eilif Muller, Srikanth Ramaswamy, Michael W. Reimann, Marwan Abdellah, Carlos Aguado Sanchez, Anastasia Ailamaki, Lidia Alonso-Nanclares, Nicolas Antille, Selim Arsever, Guy Antoine Atenekeng Kahou, Thomas K. Berger, Ahmet Bilgili, Nenad Buncic, Athanassia Chalimourda, Giuseppe Chindemi, Jean-Denis Courcol, Fabien Delalondre, Vincent Delattre, Shaul Druckmann, Raphael Dumusc, James Dynes, Stefan Eilemann, Eyal Gal, Michael Emiel Gevaert, Jean-Pierre Ghobril, Albert Gidon, Joe W. Graham, Anirudh Gupta, Valentin Haenel, Etay Hay, Thomas Heinis, Juan B. Hernando, Michael Hines, Lida Kanari, Daniel Keller, John Kenyon, Georges Khazen, Yihwa Kim, James G. King, Zoltan Kisvarday, Pramod Kumbhar, S´ebastien Lasserre, Jean-Vincent Le B´e, Bruno R. C. Magalh˜aes, Angel Merch´an-P´erez, Julie Meystre, Benjamin Roy Morrice, Jeffrey Muller, Alberto Mun˜oz-C´espedes, Shruti Muralidhar, Keerthan Muthurasa, Daniel Nachbaur, Taylor H. Newton, Max Nolte, Aleksandr Ovcharenko, Juan Palacios, Luis Pastor, Rodrigo Perin, Rajnish Ranjan, Imad Riachi, Jos´e-Rodrigo Rodr´ıguez, Juan Luis Riquelme, Christian Rössert, Konstantinos Sfyrakis, Ying Shi, Julian C. Shillcock, Gilad Silberberg, Ricardo Silva, Farhan Tauheed, Martin Telefont, Maria Toledo-Rodriguez, Thomas Tränkler, Werner Van Geit, Jafet Villafranca D´ıaz, Richard Walker, Yun Wang, Stefano M. Zaninetta, Javier DeFelipe, Sean L. Hill, Idan Segev, and Felix Schürmann. Reconstruction and simulation of neocortical microcircuitry. Cell, 163(2):456–492, 2015/10/09 2015.

3. Frances K Skinner. Cellular-based modeling of oscillatory dynamics in brain networks. Current opinion in neurobiology, 22(4):660–669, 2012.

4. Michael Hawrylycz, Costas Anastassiou, Anton Arkhipov, Jim Berg, Michael Buice, Nicholas Cain, Nathan W Gouwens, Sergey Gratiy, Ramakrishnan Iyer, Jung Hoon Lee, et al. Inferring cortical function in the mouse visual system through large-scale systems neuroscience. Proceedings of the National Academy of Sciences, 113(27):7337–7344, 2016.

5. Patricia S Churchland and Terrence J Sejnowski. Blending computational and experimental neuroscience. Nature Reviews Neuroscience, 2016.

6. Anne K Churchland and L F Abbott. Conceptual and technical advances define a key moment for theoretical neuroscience. Nat Neurosci, 19(3):348–349, 03 2016.

7. John P Cunningham and M Yu Byron. Dimensionality reduction for large-scale neural recordings. Nature neuroscience, 2014.

8. Ruben A. Tikidji-Hamburyan, Vikram Narayana, Zeki Bozkus, and Tarek A. El-Ghazawi. Software for brain network simulations: A comparative study. Frontiers in Neuroinformatics, 11:46, 2017.

9. Robert A. McDougal, Thomas M. Morse, Ted Carnevale, Luis Marenco, Rixin Wang, Michele Migliore, Perry L. Miller, Gordon M. Shepherd, and Michael L. Hines. Twenty years of modeldb and beyond: building essential modeling tools for the future of neuroscience. Journal of Computational Neuroscience, pages 1–10, 2016.

10. WW Lytton, AH Seidenstein, S Dura-Bernal, RA McDougal, F Schürmann, and ML Hines. Simulation neurotechnologies for advancing brain research: parallelizing large networks in NEURON. Neural Comput, 28:2063–2090, 2016.

11. Lealem Mulugeta, Andrew Drach, Ahmet Erdemir, CA Hunt, Marc Horner, Joy P Ku, Jerry G Myers, Jr, Rajanikanth Vadigepalli, and William W Lytton. Credibility, replicability, and reproducibility in simulation for biomedicine and clinical applications in neuroscience. Front. Neuroinform., 12:18, April 2018.

12. RA McDougal, AS Bulanova, and WW Lytton. Reproducibility in computational neuroscience models and simulations. IEEE Trans Biomed Eng, 63:2021–2035, 2016.

13. Padraig Gleeson, Sharon Crook, Robert C. Cannon, Michael L. Hines, Guy O. Billings, Matteo Farinella, Thomas M. Morse, Andrew P. Davison, Subhasis Ray, Upinder S. Bhalla, Simon R. Barnes, Yoana D. Dimitrova, and R. Angus Silver. Neuroml: A language for describing data driven models of neurons and networks with a high degree of biological detail. PLoS Comput Biol, 6(6):e1000815, 06 2010.

14. Robert C Cannon, Padraig Gleeson, Sharon Crook, Gautham Ganapathy, Boris Marin, Eugenio Piasini, and R. Angus Silver. LEMS: A language for expressing complex biological models in concise and hierarchical form and its use in underpinning NeuroML 2. Frontiers in Neuroinformatics, 8(79), 2014.

15. Robert A McDougal, Anna S Bulanova, and William W Lytton. Reproducibility in computational neuroscience models and simulations. IEEE Transactions on Biomedical Engineering, 63(10):2021–2035, 2016.

16. Padraig Gleeson, Volker Steuber, and R Angus Silver. neuroconstruct: a tool for modeling networks of neurons in 3d space. Neuron, 54(2):219–235, 2007.

17. Andrew Davison, Daniel Brüderle, Jochen Eppler, Jens Kremkow, Eilif Muller, Dejan Pecevski, Laurent Perrinet, and Pierre Yger. Pynn: a common interface for neuronal network simulators. Frontiers in Neuroinformatics, 2:11, 2009.

18. James Bednar. Topographica: building and analyzing map-level simulations from python, c/c++, matlab, nest, or neuron components. Frontiers in Neuroinformatics, 3:8, 2009.

19. Sergey G. Aleksin, Kaiyu Zheng, Dmitri A. Rusakov, and Leonid P. Savtchenko. Arachne: A neural-neuroglial network builder with remotely controlled parallel computing. PLOS Computational Biology, 13(3):1–14, 03 2017.

20. Sergey L. Gratiy, Yazan N. Billeh, Kael Dai, Catalin Mitelut, David Feng, Nathan W. Gouwens, Nicholas Cain, Christof Koch, Costas A. Anastassiou, and Anton Arkhipov. Bionet: A python interface to neuron for modeling large-scale networks. PLOS ONE, 13(8):1–18, 08 2018.

21. Robert McDougal, Michael Hines, and William Lytton. Reaction-diffusion in the neuron simulator. Frontiers in Neuroinformatics, 7:28, 2013.

22. Adam J. H. Newton, Robert A. McDougal, Michael L. Hines, and William W. Lytton. Using neuron for reaction-diffusion modeling of extracellular dynamics. Frontiers in Neuroinformatics, 12:41, 2018.

23. S Dura-Bernal, SA Neymotin, BA Suter, GMG Shepherd, and WW Lytton. Long-range inputs and h-current regulate different modes of operation in a multiscale model of mouse m1 microcircuits. bioRxiv, (201707), 2018.

24. Cecilia Romaro, Fernando Araujo Najman, Salvador Dura-Bernal, and Antonio Carlos Roque. Implementation of the potjans-diesmann cortical microcircuit model in netpyne/neuron with rescaling option. In 27th Annual Computational Neuroscience Meeting, CNS*18, 2018.

25. William Lytton, Jing Xuan Limb, Salvador Dura-Bernal, and George J. Augustine. Computer models of claustrum subnetworks. In Brian N. Mathur, David Reser, and Jared B. Smith, editors, Conference Proceedings: 3rd Annual Society for Claustrum Research Meeting, Claustrum, volume 2:1, page 1349859, 2017.

26. Samuel A Neymotin, Salvador Dura-Bernal, Peter Lakatos, Terence David Sanger, and William Lytton. Multitarget multiscale simulation for pharmacological treatment of dystonia in motor cortex. Frontiers in Pharmacology, 7(157), 2016.

27. Stephanie R Jones, Dominique L Pritchett, Michael A Sikora, Steven M Stufflebeam, Matti Hämäläinen, and Christopher I Moore. Quantitative analysis and biophysically realistic neural modeling of the meg mu rhythm: rhythmogenesis and modulation of sensory-evoked responses. Journal of neurophysiology, 102(6):3554–3572, 2009.

28. S. A. Neymotin, D. S. Daniels, N. Peled, R. A. McDougal, N. T. Carnevale, C. I. Moore, S. Dura-Bernal, ML Hines, and S Jones. Human neocortical neurosolver (hnn). doi:10.5281/ zenodo.1446517, 2018.

29. Padraig Gleeson, Matteo Cantarelli, Boris Marin, Adrian Quintana, Matt Earnshaw, Eugenio Piasini, Justas Birgiolas, Robert C Cannon, N Alex Cayco-Gajic, Sharon Crook, et al. Open source brain: a collaborative resource for visualizing, analyzing, simulating and developing standardized models of neurons and circuits. bioRxiv, page 229484, 2018.

30. Subhashini Sivagnanam, Amit Majumdar, Kenneth Yoshimoto, Vadim Astakhov, Anita Bandrowski, Maryann E Martone, and Nicholas T Carnevale. Introducing the neuroscience gateway. In IWSG, 2013.

31. WW Lytton and M Stewart. Rule-based firing for network simulations. Neurocomputing, 69:1160–1164, 2006.

32. Richard Naud, Nicolas Marcille, Claudia Clopath, and Wulfram Gerstner. Firing patterns in the adaptive exponential integrate-and-fire model. Biol. Cybern., 99(4-5):335–347, 2008.

33. EM Izhikevich. Simple Model of Spiking Neurons. IEEE Trans Neural Networks, 14:1569–1572, 2003.

34. ML Hines and NT Carnevale. Expanding NEURON’s repertoire of mechanisms with NMODL. Neural Computation, 12:995–1007, 2000.

35. SA Neymotin, RA McDougal, AS Bulanova, M Zeki, P Lakatos, D Terman, ML Hines, and WW Lytton. Calcium regulation of hcn channels supports persistent activity in a multiscale model of neocortex. Neuroscience, 316:344–366, 2016.

36. S.L. Angulo, R. Orman, S.A. Neymotin, L. Liu, L. Buitrago, E. Cepeda-Prado, D. Stefanov, W.W. Lytton, M. Stewart, S.A. Small, K.E. Duff, and H. Moreno. Tau and amyloid-related pathologies in the entorhinal cortex have divergent effects in the hippocampal circuit. Neurobiology of Disease, 108(Supplement C): 261–276, 2017.

37. Leopoldo Petreanu, Tianyi Mao, Scott M Sternson, and Karel Svoboda. The subcellular organization of neocortical excitatory connections. Nature, 457(7233):1142–1145, 2009.

38. Charles T. Anderson, Patrick L. Sheets, Taro Kiritani, and Gordon M. G. Shepherd. Sublayer-specific microcircuits of corticospinal and corticostriatal neurons in motor cortex. Nature neuroscience, 13(6):739–44, June 2010.

39. Benjamin A Suter and Gordon MG Shepherd. Reciprocal interareal connections to corticospinal neurons in mouse m1 and s2. The Journal of Neuroscience, 35(7):2959–2974, 2015.

40. Bryan M Hooks, Tianyi Mao, Diego A Gutnisky, Naoki Yamawaki, Karel Svoboda, and Gordon M G Shepherd. Organization of cortical and thalamic input to pyramidal neurons in mouse motor cortex. J Neurosci, 33(2):748–760, Jan 2013.

41. Marianne J Bezaire, Ivan Raikov, Kelly Burk, Dhrumil Vyas, and Ivan Soltesz. Interneuronal mechanisms of hippocampal theta oscillation in a full-scale model of the rodent ca1 circuit. eLife, 5:e18566, 2016.

42. M Hereld, RL Stevens, J Teller, and W van Drongelen. Large Neural Simulations on Large Parallel Computers. Int. J. for Bioelectromagnetism, 7:44–46, 2005.

43. Michael Hines, Sameer Kumar, and Felix Schürmann. Comparison of neuronal spike exchange methods on a blue gene/p supercomputer. Frontiers in Computational Neuroscience, 5(49), 2011.

44. Michael L Hines, Hubert Eichner, and Felix Schürmann. Neuron splitting in compute-bound parallel network simulations enables runtime scaling with twice as many processors. Journal of computational neuroscience, 25(1):203–210, 2008.

45. M Migliore, C Cannia, WW Lytton, and ML Hines. Parallel network simulations with NEURON. J. Computational Neuroscience, 6:119–129, 2006.

46. John Towns, Timothy Cockerill, Maytal Dahan, Ian Foster, Kelly Gaither, Andrew Grimshaw, Victor Hazlewood, Scott Lathrop, Dave Lifka, Gregory D Peterson, et al. Xsede: accelerating scientific discovery. Computing in Science & Engineering, 16(5):62–74, 2014.

47. Katrin Amunts, Christoph Ebell, Jeff Muller, Martin Telefont, Alois Knoll, and Thomas Lippert. The human brain project: Creating a european research infrastructure to decode the human brain. Neuron, 92(3):574–581, 2017/04/17 2017.

48. Dorian Krause and Philipp Thörnig. Jureca: Modular supercomputer at jülich supercomputing centre. Journal of large-scale research facilities JLSRF, 4:132, 2018.

49. Michele Migliore, C Cannia, William W Lytton, Henry Markram, and Michael L Hines. Parallel network simulations with neuron. Journal of computational neuroscience, 21(2):119–129, 2006.

50. Tobias C. Potjans and Markus Diesmann. The cell-type specific cortical microcircuit: Relating structure and activity in a full-scale spiking network model. Cerebral Cortex, 24(3):785–806, 2014.

51. Thomas Kreuz, Mario Mulansky, and Nebojsa Bozanic. Spiky: A graphical user interface for monitoring spike train synchrony. Journal of Neurophysiology, 113(9):3432–3445, 2015.

52. H Lind´en, E Hagen, S Leski, ES Norheim, KH Pettersen, and GT Einevoll. LFPy: a tool for biophysical simulation of extracellular potentials generated by detailed model neurons. Front Neuroinform, 7:41, 2013.

53. Harilal Parasuram, Bipin Nair, Egidio D’Angelo, Michael Hines, Giovanni Naldi, and Shyam Diwakar. Computational modeling of single neuron extracellular electric potentials and network local field potentials using lfpsim. Frontiers in Computational Neuroscience, 10:65, 2016.

54. György Buzs´aki, Costas A Anastassiou, and Christof Koch. The origin of extracellular fields and currents—eeg, ecog, lfp and spikes. Nature reviews neuroscience, 13(6):407, 2012.

55. Dan Goodman and Romain Brette. Brian: a simulator for spiking neural networks in python. Frontiers in Neuroinformatics, 2:5, 2008.

56. JM Bower and D Beeman. The book of Genesis: exploring realistic neural models with the GEneral NEural SImulation System. 1998. Science and Business Media, Springer, New York, 2012.

57. Subhasis Ray and Upinder Bhalla. Pymoose: interoperable scripting in python for moose. Frontiers in Neuroinformatics, 2:6, 2008.

58. Gerald M Edelman and Joseph A Gally. Degeneracy and complexity in biological systems. Proc Nat Acad Sci, 98:13763–13768, 2001.

59. AA Prinz, D Bucher, and E Marder. Similar network activity from disparate circuit parameters. Nat Neurosci, 7:1345–1352, 2004.

60. SA Neymotin, S Dura-Bernal, P Lakatos, TD Sanger, and WW Lytton. Multitarget multiscale simulation for pharmacological treatment of dystonia in motor cortex. Front Pharmacol, 7:157, 2016.

61. Samuel A Neymotin, Heekyung Lee, Eunhye Park, Andre A Fenton, and William W Lytton. Emergence of physiological oscillation frequencies in a computer model of neocortex. Front Comput Neurosci, 5:19, 2011.

62. Andrew T Knox, Tracy Glauser, Jeffrey Tenney, William W Lytton, and Katherine Holland. Modeling pathogenesis and treatment response in childhood absence epilepsy. Epilepsia, 59(1):135–145, 2018.

63. W Van Geit, Erik De Schutter, and Pablo Achard. Automated neuron model optimization techniques: a review. Biological cybernetics, 99(4-5):241–251, 2008.

64. Carmen G Moles, Pedro Mendes, and Julio R Banga. Parameter estimation in biochemical pathways: a comparison of global optimization methods. Genome research, 13(11):2467–2474, 2003.

65. S. Dura-Bernal, S. A. Neymotin, C. C. Kerr, S. Sivagnanam, A. Majumdar, J. T. Francis, and W. W. Lytton. Evolutionary algorithm optimization of biological learning parameters in a biomimetic neuroprosthesis. IBM Journal of Research and Development, 61(2/3):6:1–6:14, March 2017.

66. SA Neymotin, BA Suter, S Dura-Bernal, GM Shepherd, M Migliore, and WW Lytton. Optimizing computer models of corticospinal neurons to replicate in vitro dynamics. J Neurophysiol, 117:148–162, 2017.

67. Kristofor David Carlson, Jayram M Nageswaran, Nikil Dutt, and Jeffrey L Krichmar. An efficient automated parameter tuning framework for spiking neural networks. Frontiers in Neuroscience, 8(10), 2014.

68. Timothy H Rumbell, Danel Dragulji´c, Aniruddha Yadav, Patrick R Hof, Jennifer I Luebke, and Christina M Weaver. Automated evolutionary optimization of ion channel conductances and kinetics in models of young and aged rhesus monkey pyramidal neurons. Journal of Computational Neuroscience, 41(1):65–90, 2016.

69. Pablo Achard and Erik De Schutter. Complex parameter landscape for a complex neuron model. PLoS Comput Biol, 2(7):e94, 2006.

70. Nathan W Gouwens, Jim Berg, David Feng, Staci A Sorensen, Hongkui Zeng, Michael J Hawrylycz, Christof Koch, and Anton Arkhipov. Systematic generation of biophysically detailed models for diverse cortical neuron types. Nature Communications, 9(1):710, 2018.

71. Robert A McDougal, Thomas M Morse, Michael L Hines, and Gordon M Shepherd. ModelView for ModelDB: Online presentation of model structure. Neuroinformatics, 13(4):459–470, October 2015.

72. Benjamin A Suter, Michele Migliore, and Gordon MG Shepherd. Intrinsic electrophysiology of mouse corticospinal neurons: a class-specific triad of spike-related properties. Cerebral Cortex, 23(8):1965–1977, 2013.

73. Naoki Yamawaki, Katharine Borges, Benjamin A Suter, Kenneth D Harris, and Gordon MG Shepherd. A genuine layer 4 in motor cortex with prototypical synaptic circuit connectivity. Elife, 3:e05422, 2015.

74. Naoki Yamawaki and Gordon M G Shepherd. Synaptic circuit organization of motor corticothalamic neurons. The Journal of Neuroscience, 35(5):2293–2307, 2015.

75. Kenneth D Harris and Gordon MG Shepherd. The neocortical circuit: themes and variations. Nature Neuroscience, 18(2):170–181, 2015.

76. Patrick L Sheets, Benjamin A Suter, Taro Kiritani, C Savio Chan, D James Surmeier, and Gordon MG Shepherd. Corticospinal-specific hcn expression in mouse motor cortex: Ih-dependent synaptic integration as a candidate microcircuit mechanism involved in motor control. Journal of neurophysiology, 106(5):2216–2231, 2011.

77. Nicholas Weiler, Lydia Wood, Jianing Yu, Sara A Solla, and Gordon M G Shepherd. Top-down laminar organization of the excitatory network in motor cortex. Nat Neurosci, 11(3):360–366, Mar 2008.

78. Taro Kiritani, Ian R. Wickersham, H. Sebastian Seung, and Gordon M. G. Shepherd. Hierarchical connectivity and connection-specific dynamics in the corticospinal–corticostriatal microcircuit in mouse motor cortex. The Journal of Neuroscience, 32(14):4992–5001, 2012.

79. Alfonso J Apicella, Ian R Wickersham, H Sebastian Seung, and Gordon MG Shepherd. Laminarly orthogonal excitation of fast-spiking and low-threshold-spiking interneurons in mouse motor cortex. The Journal of Neuroscience, 32(20):7021–7033, 2012.

80. Julian Tejada, Norberto Garcia-Cairasco, and Antonio C Roque. Combined role of seizureinduced dendritic morphology alterations and spine loss in newborn granule cells with mossy fiber sprouting on the hyperexcitability of a computer model of the dentate gyrus. PLoS computational biology, 10(5):e1003601, 2014.

81. Facundo Rodriguez. Dentate gyrus network model. In 27th Annual Computational Neuroscience Meeting, CNS*18, 2018.

82. Vassilis Cutsuridis, Stuart Cobb, and Bruce P Graham. Encoding and retrieval in a model of the hippocampal ca1 microcircuit. Hippocampus, 20(3):423–446, 2010.

83. Angeles Tepper, Adam Sugi, William Lytton, and Salvador Dura-Bernal. Implementation of ca1 microcircuits model in netpyne and exploration of the effect of neuronal/synaptic loss on memory recall. In 27th Annual Computational Neuroscience Meeting, CNS*18, 2018.

84. Salvador Dura-Bernal, Kan Li, Samuel A Neymotin, Joseph T Francis, Jose C Principe, and William W Lytton. Restoring behavior via inverse neurocontroller in a lesioned cortical spiking model driving a virtual arm. Frontiers in Neuroscience, 10(28), 2016.

85. Alexander O. Komendantov and Giorgio A. Ascoli. Dendritic excitability and neuronal morphology as determinants of synaptic efficacy. Journal of Neurophysiology, 101(4):1847–1866, 2009.

86. Yoshiyuki Kubota, Satoru Kondo, Masaki Nomura, Sayuri Hatada, Noboru Yamaguchi, Alsayed A Mohamed, Fuyuki Karube, Joachim Lübke, and Yasuo Kawaguchi. Functional effects of distinct innervation styles of pyramidal cells by fast spiking cortical interneurons. eLife, 4:e07919, jul 2015.

87. Werner Van Geit, Michael Gevaert, Giuseppe Chindemi, Christian Rössert, Jean-Denis Courcol, Eilif Muller, Felix Schürmann, Idan Segev, and Henry Markram. Bluepyopt: Leveraging open source software and cloud infrastructure to optimise model parameters in neuroscience. arXiv preprint arXiv:1603.00500, 2016.

88. P´eter Friedrich, Michael Vella, Attila I Guly´as, Tam´as F Freund, and Szabolcs K´ali. A flexible, interactive software tool for fitting the parameters of neuronal models. Frontiers in Neuroinformatics, 8(63), 2014.

89. Robert Meyer and Klaus Obermayer. pypet: A python toolkit for data management of parameter explorations. Frontiers in Neuroinformatics, 10:38, 2016.

90. 2017.

91. Carl-Magnus Svensson, Stephen Coombes, and JonathanWestley Peirce. Using evolutionary algorithms for fitting high-dimensional models to neuronal data. Neuroinformatics, 10(2):199–218, 2012.

92. Maxwell A Sherman, Shane Lee, Robert Law, Saskia Haegens, Catherine A Thorn, Matti S Hämäläinen, Christopher I Moore, and Stephanie R Jones. Neural mechanisms of transient neocortical beta rhythms: Converging evidence from humans, computational modeling, monkeys, and mice. Proceedings of the National Academy of Sciences, page 201604135, 2016.

93. WW Lytton. Computer modelling of epilepsy. Nat Rev Neurosci, 9:626–637, 2008.

94. John K Salmon, Mark A Moraes, Ron O Dror, and David E Shaw. Parallel random numbers: as easy as 1, 2, 3. In Proceedings of 2011 International Conference for High Performance Computing, Networking, Storage and Analysis, page 16. ACM, 2011.

95. Matteo Cantarelli, Boris Marin, Adrian Quintana, Matt Earnshaw, Padraig Gleeson, Salvador Dura-Bernal, R Angus Silver, and Giovanni Idili. Geppetto: a reusable modular open platform for exploring neuroscience data and models. Phil. Trans. R. Soc. B, 373(1758):20170380, 2018.

